# Improved single-molecule localization precision in astigmatism-based 3D superresolution imaging using weighted likelihood estimation

**DOI:** 10.1101/304816

**Authors:** Christopher H. Bohrer, Xinxing Yang, Zhixin Lyu, Shih-Chin Wang, Jie Xiao

## Abstract

Astigmatism-based superresolution microscopy is widely used to determine the position of individual fluorescent emitters in three-dimensions (3D) with subdiffraction-limited resolutions. This point spread function (PSF) engineering technique utilizes a cylindrical lens to modify the shape of the PSF and break its symmetry above and below the focal plane. The resulting asymmetric PSFs at different z-positions for single emitters are fit with an elliptical 2D-Gaussian function to extract the widths along two principle x- and y-axes, which are then compared with a pre-measured calibration function to determine its z-position. While conceptually simple and easy to implement, in practice, distorted PSFs due to an imperfect optical system often compromise the localization precision; and it is laborious to optimize a multi-purpose optical system. Here we present a methodology that is independent of obtaining a perfect PSF and enhances the localization precision along the z-axis. By utilizing multiple calibration images of fluorescent beads at varying z-planes and characterizing experimentally measured background distributions, we numerically approximated the probability of observing a certain signal in a given pixel from a single emitter at a particular z-plane. We then used a weighted maximum likelihood estimator (WLE) to determine the 3D-position of the emitter. We demonstrate that this approach enhances z-axis localization precision in all conditions we tested, in particular when the PSFs deviate from a standard 2D Gaussian model.

## Introduction

Single molecule localization microscopy (SMLM) relies on the temporal isolation of individual fluorescence emitters to determine the spatial localizations of individual molecules with high precision(1–4). SMLM has been widely used in biology to address the spatial organizations and structural dimensions of sub-cellular structures at a resolution ~ 10-fold better than the diffraction limit of conventional fluorescence light microscopy(5–8). Localization precision, the error in determining the spatial coordinates of a single emitter, is a critical parameter of SMLM. Together with sample labeling density(9), localization precision determines the upper bound of achievable spatial resolution(10). Two-dimensional (2D) SMLM methods can reach a lateral localization precision of 10-40 nm along the *x* and *y* dimensions in the focal plane by fitting the single emitter’s image to a point spread function (PSF) model. The PSF is usually approximated by using a symmetric 2D-Gaussian function in most algorithms(1–3, 11). However, in practice, the true PSF of a given imperfect optical system can deviate significantly from the symmetric 2D-Gaussian function (12).

Recent developments in SMLM have allowed the coordinate of an emitter along the third dimension, the z-axis of the optical path, to be determined with a precision in the range of 15 – 100 nm. There are two major approaches(13, 14): interferometry-based methods such as interferometric photoactivated localization microscopy (iPALM (15)), and PSF-engineering/extension-based methods such as astigmatism (AS)(16), double-helix (DH)(17), and bi/multi-focal plane (BP) microscopy(18). Among these methods, iPALM uses the interference of the same photon emitted from an emitter to reach the highest localization precision along the z-dimension at ~ 15 nm. However, the relatively narrow observation depth (<750 nm above the coverslip(19)) and complex microscopy setup have limited its broad applications in biology. Multifocal plane or PSF-engineering methods determine the z-position of single emitters by comparing the image of a single emitter with calibrated PSFs at different z-planes, and in general, can reach a z-resolution in the range of 40 – 80 nm. Although the z-axis resolution is about 2-3 times worse than the lateral resolution, these PSF-engineering methods are easy and of low cost to implement. In particular, astigmatism-based 3D-SMLM only requires adding a cylindrical lens in the emission pathway, and hence has seen broad applications within the biological imaging community (16, 17, 20).

In an ideal astigmatism-based optical system, the PSF of a freely rotating single fluorescence emitter can be mimicked by a 2D-Gaussian function (16)

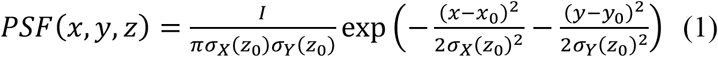

Where the *x*_0_, *y*_0_, and *z*_0_ are the true spatial coordinates of the emitter, and σ_*X(Z_0_)*_ and σ_*Y(Z_0_)*_ present the widths of the Gaussian function along two perpendicular *x* and *y*-axes at *z*_0_. Here, for simplicity, the *x* and *y*-axes represent the principle axes of the cylindrical lens respectively. In all astigmatism-based 3D-SMLM imaging, a calibration curve describing the correlation between the astigmatism of an emitter’s PSF and its *z*-position is first established by imaging a fluorescent bead smaller than the diffraction limit at predefined *z*-planes, and subsequently extracting the astigmatic widths of the emitter’s PSF using equation 1. The extracted widths are fitted as a function of the corresponding *z*-positions using a phenomenological model such as the defocusing function(16) or the quadratic function(21, 22). By comparing experimentally measured widths of a single emitter with the calibration curves, one can obtain the *z* position of the emitter with a *z*-axis localization precision of 40~80 nm under the condition of a nearly perfect optical setup (so that the PSF can be approximated by equation 1) (16, 17, 20). However, in practice, it is laborious to perfect the optics each time on a multi-purpose microscope, even for experienced users. As such, imperfect experimental optical setups lead to distorted PSFs and consequently large deviations of measured calibration curves away from these commonly used models, introducing significant uncertainties in determining an emitter’s *z*-position (Fig. S1).

To reduce the discrepancy between experimentally measured calibration curves and the fitting models, Sauer’s group used B-spline to interpolate the calibration curves (Fig. S1), which was able to obtain higher accuracy and flexibility than the original fitting functions under various experimental conditions(23). Additionally, Shaevitz *et al.* used a Bayesian interference method to measure the probability distribution of astigmatic PSF widths at different *z*-positions(24). Nevertheless, these methods still assumed that each emitter’s PSF could be approximated by an elliptical 2D Gaussian function, which doesn’t necessarily hold true in an imperfect optical system (Figure 1A). For instance, the maximum intensity position of the PSF can shift, or ‘wobble,’ due to coverslip-tilt and non-rotational symmetric aberration of an individual objective or other components in microscope(13). Additionally, spherical and other aberrations can distort the PSF shape, which introduces bias and compromise the localization precision in the *z*-position (as well as in *x-y*) (12) (Fig. 1A).

**Figure 1.**
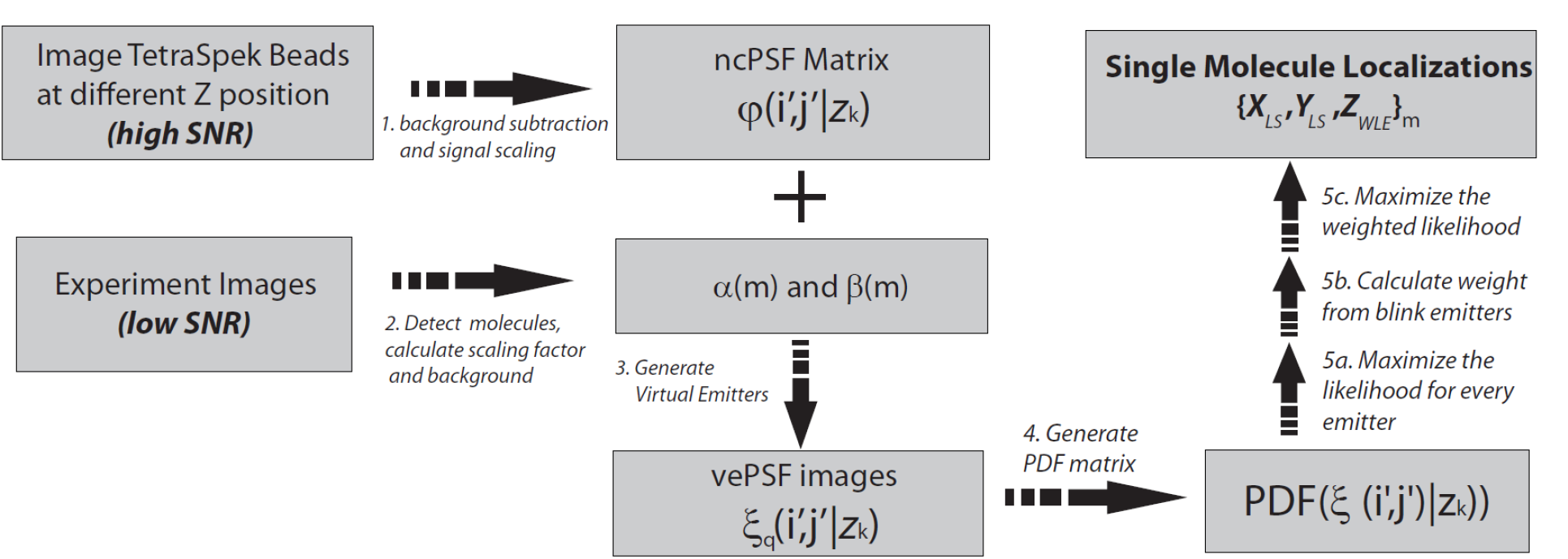
Schematics of the WLE workflow. Experimentally measured bead images at different z-planes and real emitter images were used for localization.

An analytical description of the PSF, under specific conditions, can be retrieved from the pupil function of the imaging setup and has been shown to improve z-axis resolution(25, 26). The pupil function is then interpolated or decomposed in Zernike polynomials to calculate the PSF at different *z*-positions. The 3D phase retrieval (PR) method has been implemented in BP and DH microscopy to approximate the pupil function(25, 27). However, these PDF-retrieval methods are tedious to implement and require a thorough understanding of the optical setup.

In this work, we introduce a different approach using the experimentally measured PSF and a numerically weighted maximum likelihood estimation (WLE) to improve the *z*-axis localization precision of single emitters in astigmatism-based SMLM. Our method is based on the principle that the PSF of an emitter at different *z*-positions can be characterized as an experimentally measured image independent of any a priori model assumptions such as an elliptical Gaussian model. For each experimentally measured image of an emitter, our method numerically determines the probability for each pixel to have a particular signal level given its *z*-position using the image of a calibration bead at each *z*-plane with an experimentally characterized background noise distribution. We then weight the importance of the pixels of an emitter by conducting a phase space search to minimize the calculated z-distances between repeated localizations from the same emitters. We verified that by maximizing the weighted likelihood, we could reach a comparable or higher localization precision when compared to the B-spline based traditional Least Square (LS) fitting methodologies for 2D Gaussian PSFs, independent of the optical setup. It shows the highest improvement when the experimentally measured PSF deviates significantly from the standard elliptical 2D Gaussian model. Thus, the WLE approach alleviates the practical concerns in perfecting the optical alignment and enables improved *z*-axis localization in astigmatism-based 3D superresolution imaging.

## Method and Results

### Principle and workflow of WLE

To estimate an emitter’s coordinates, least square fitting (LS) and maximum likelihood estimator (MLE) based on a ‘known’ PSF, are the most commonly used methods. LS is computationally faster than MLE and has produced comparable precision in experiments where high photon counts are achievable(28). In SMLM fitting algorithms, LS fitting is often chosen over MLE to increase the computational speed and simplify the fitting process. However, when the signal to noise ratio (SNR) is low, LS leads to a compromised localization precision. MLE is more computationally intensive compared to LS, but with a ‘correct’ PSF function and a ‘correct’ noise model (11), one can theoretically approach the upper bound of the estimation precision, the Crame-Rao limitation(29).

Our approach, termed weighted maximum likelihood estimation (WLE), determines a single emitter’s *z*-position by maximizing the weighted likelihood of having a particular signal for each pixel at a particular z-position according to an experimentally determined probability density matrix (PDM). WLE is very similar to the MLE approach, but it assigns the information from different sources, different pixels in this case, varying degrees of (weighted) importance. The PDM was determined by convolving the numerically calibrated point spread function (ncPSF) with scaled photon noise and the background noise distribution. In essence, WLE finds a single emitter’s z-position by numerically matching its experimentally measured image with that of a calibration bead at a known *z*-position considering the intensity and background distribution of each pixel. Below, we describe the five main steps to implement the WLE algorithm (Fig. 1).

In the first step, we estimated the ncPSF by imaging a bright fluorescent bead (100 nm in diameter) at a series of evenly spaced, 10-nm apart, *z*-planes using a piezo-stage. As a point source, the background-free averaged and normalized image represents the ncPSF of the optical system. To increase the signal to noise ratio, we took multiple (*N*) images at each z-plane. We then computed the ncPSF, *φ_ij_*(*k*), for each *k*^th^ z-plane as:

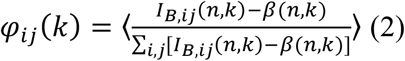

where *I_B,ij_*(*n, k*) is the intensity of the pixel at row *i* and column *j* in the *n*^th^ bead image (21 by 21 pixels in our example and can be varied) at the *k*^th^ z-plane, and *β*(*n, k*) is the background intensity calculated using averaged intensity values of pixels furthest away from the bead center. The mean background intensity *β*(*n, k*) was subtracted from *I_B,ij_*(*n, k*) to obtain the true signal intensity at each pixel, which was further normalized by the integrated signal intensity of the image after background subtraction. The final ncPSF was obtained by averaging over all *N* images of the bead for the *k*^th^ *z*-plane. This estimation of ncPSF is sufficient because of the negligible noise level due to the high signal from fluorescent beads and averaging over *N* images allows us to approximate the final background of the mean image as zero (Supporting Material).

In the second step, we obtained images containing single-molecule emitters from simulation or experimentally measured astigmatism-based imaging (Supporting Material). A wavelet-filter based algorithm(30) was applied to identify and crop out the local maxima into regions of individual emitters with the same size as ncPSF. We then calculated the background noise distribution, *β*, using the peripheral pixels in the cropped images. Here, we assumed that the noise distribution was identical among the pixels in all cropped images, which can be further adjusted depending upon whether the background is uniform among pixels. To obtain the PDM, we then determined the scaling factor *α*(*m*) with the following:

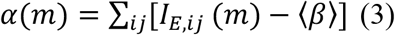

where *I_E,ij_*(*m*) is the intensity of each pixel for the cropped emitter *m*. Therefore, *α*(*m*) is proportional to the total photon number from each single emitter. Here we assumed that the distribution of the scaling factors among different emitters was independent of their *z*-positions. We validated this assumption using experimental data (Fig. S2, Supporting Material).

In the third step, we determined the PDM, which allowed us to calculate the weighted likelihood. We utilized the ncPSF, {*φ_ij_*(*k*), *k* = 1,2,…} estimated in Step 1 to generate *q* calibration emitters at each z-plane *k*, {*ξ_ij_*(*q, k*)}, incorporating the previously determined background noise *β* and scaling factors *α*. Note here that *β* and *α* are randomly sampled variables from their corresponding distributions (Supporting Material). To generate the calibration emitters, we first simulated a signal for each pixel using the ncPSF multiplied by the scaling factor *α* with Poisson noise (photon noise) in each pixel: *Poisson*[*φ_ij_*(*k*) × *α*]. Here we assumed Poisson noise for simplicity, but other circumstances could be accommodated. The background noise of each pixel was added by randomly sampling the noise distribution *β*. For simplicity and computational ease, we generated 4000 calibration emitters at each z-plane with varying *α* and background noise. In principle, one could instead generate the calibration emitters at each *z*-plane for each specific *α* and incorporate the specific background noise distribution of an emitter’s cropped pixels, which will likely increase *z*-axis localization precision even further. Here the experimentally calibrated emitter image, which we named ecPSF, is:

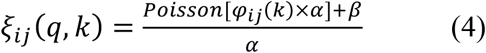

Here *α*, the scaling factor, is randomly sampled from its distribution and has the same value in both the denominator and numerator of the equation. We then linearly shifted the centroids of the ecPSFs (estimated using a 2D-Gaussian fitting) so that the “centers” of all the adjusted PSF’s were aligned at each *z*-plane.

Next, we approximated the probability density distribution to observe the signal, 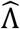 at pixel (*i*’, *j*’) for the *k*^th^ *z*-plane with 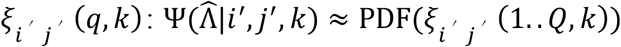, where PDF is the approximated probability density function for the term within the parentheses and *Q* is the total number of experimental calibration emitters for that *z*-plane (Supporting Material). Here 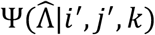 is what we referred to as the PDM, the probability density matrix, which we then used below to calculate the likelihood.

In the last step, for each single emitter image, we normalized the intensity to the total intensity of each image after background subtraction and determined the centroid as in the previous steps. This procedure resulted in the adjusted signal ∧*_m_*(*i’,j’*) for the *m*^th^ emitter. We then determined the optimal *z*-position for the *m*^th^ emitter by maximizing the following: *L*(*k*) = Σ ω(*i’,j’*) × log (Ψ(∧_*m*_(i’, j’)|i’, j’, k)) for the *k*^th^ z-plane. Here the elements of ω contain the weighted importance for each pixel (Supporting Material, Fig. S3). We determined the weights that resulted in the best resolution by performing a phase space search, adjusting each element of ω to minimize the distance between repeated localizations of the same emitters (Supporting Material, Fig. S4). (We provide a user guide, code and example data allowing one to implement and understand the inner workings of WLE (https://github.com/XiaoLabJHU/WLE).)

### Validation of WLE

To validate the WLE algorithm, we imaged TetraSpeck™ fluorescence beads on a coverslip scanning 100 z-planes at 10-nm intervals. For each z-plane, we acquired 200 images and observed minimal photobleaching. For astigmatism-based 3D SMLM imaging, it is necessary to adjust the objective correction collar and the position of the cylindrical lens to minimize the spherical aberration caused by refractive index mismatch and the cylindrical lens itself. We mimicked these adjustments and obtained three sets of calibration PSF images at different settings (PSF I, II and III, Fig. 2A). All these experimentally measured PSFs deviated from the perfect 2D-Gaussian function (Fig. 2B). Nevertheless, as a comparison, we fit these PSFs using Equation 1 to obtain the centroid positions (*x*_0_, *y*_0_) and the astigmatic PSF widths, *σ*_*X*(*z*_0_)_ and *σ*_*Y*(*z*_0_)_, at various z-planes (Fig. S1A). As shown in Fig. S1A, although the B-spline function fit the correlation between the widths and *z*-plane significantly better than a quadratic function, the correlation shape and the corresponding errors varied dramatically for the three different conditions, indicating high levels of uncertainty introduced by LS-based 2D-Gaussian fitting due to distorted PSF shapes.

**Figure 2.**
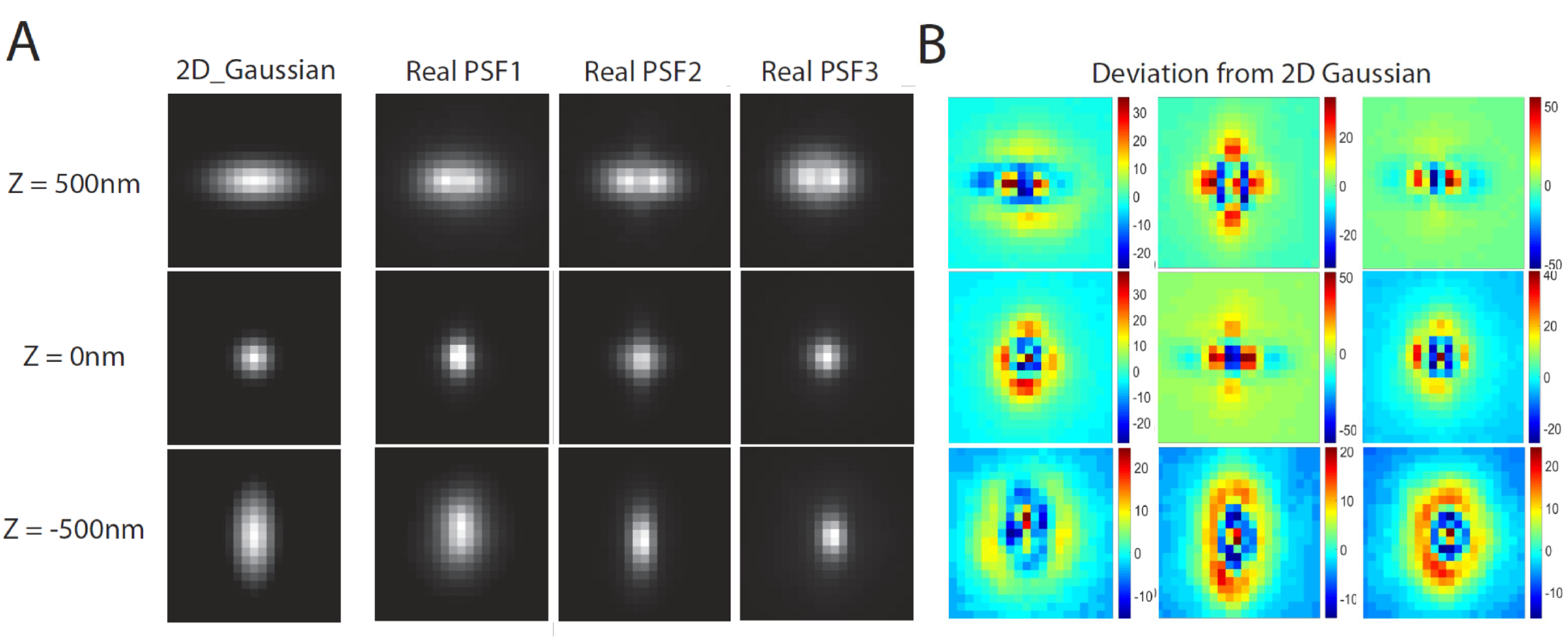
Deviation of experimentally measured ncPSF from a 2D-Gaussian model. (A) Simulated 2D-Gaussian PSF image (first column) and TetraSpeck™ beads images (experimental ncPSF) at different z-planes (−500, 0, 500 nm) with three different optical setups. PSF2 is adjusted from PSF1 by shifting the cylindrical lens position along the optical axis while PSF3 is by changing the correction collar position of the objective. (B) Numerical deviation of experimental ncPSFs from the corresponding best 2D-Gaussian fittings.

To evaluate the performance of WLE in comparison with LS-based B-spline and quadratic fitting methods, we simulated single emitters for each of these experimental PSFs at various *z*-planes with different signal to noise ratios (SNRs) (Fig. 3, Table S1 and Supporting Material). For LS-based B-spline and quadratic fitting methods, we fitted images of single emitters from highest SNR data (Table S1) with Equation 1 to obtain the z-positions using different calibration functions (B-spline or quadratic (Fig. 3). Here we defined the error, or the localization precision, as the mean absolute distance of all emitters from their true locations under each condition. As shown in Fig. 3, for all three different optical settings and at different SNR, the quadratic function (purple) performed most poorly and reached a plateau of error at ~ 40 to 60 nm for PSF I and III. The B-spline method was significantly better than the quadratic method and was able to reach a maximum resolution of ~10 nm with the highest SNR for all three PSFs, suggesting that it is more reliable and adaptive in fitting the calibration curve compared to the quadratic function, as shown previously(23).

**Figure 3.**
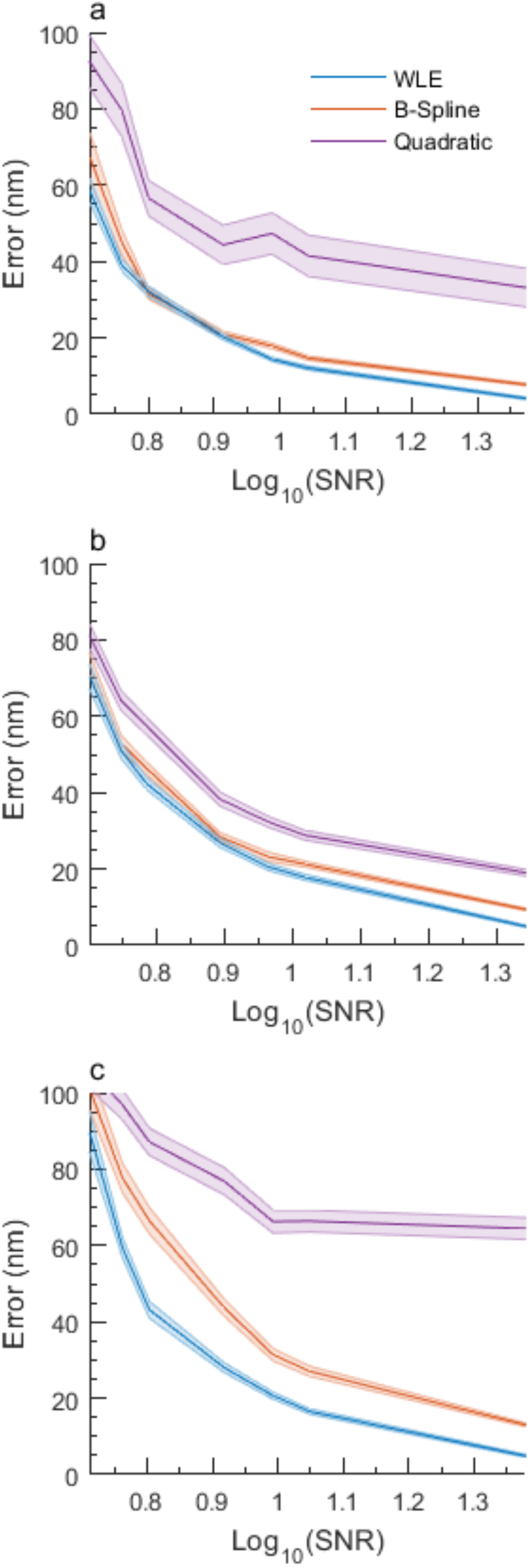
Average localization precision of LS-fitting (purple for quadratic and orange for B-spline) and WLE (blue) using synthetic images generated from experimental ncPSFs (Figure 2) at a series of signal to noise ratio (SNR).

For WLE, we determined the z-position of each simulated emitter by comparing its corresponding image with the bead-generated ncPSF and maximizing the weighted likelihood. For PSFs I and II, we found that WLE resulted in similar localization precisions compared to B-spline when SNRs were low but showed more significant improvement than B-spline when the SNR was high. For PSF III, we observed the greatest improvement (~1.5-fold) by WLE compared to B-spline, reaching a localization precision of < 10 nm and surpassing all other methodologies for every SNR we tested. These results illustrated that WLE consistently performed equally well or better than the best-performing LS-based B-spline method under all tested conditions.

### WLE improved z-axis localization recision independent of PSF shape

Next, we reasoned that the varied levels of improvement of WLE over B-spline (Fig. 3A to C) could be due to the deviation of the experimentally measured PSFs from an ideal 2D Gaussian function — the larger the deviation, the better WLE outperforms the LS methods. Therefore, we quantified the deviation of a PSF from an ideal elliptical 2D Gaussian as the mean of the absolute difference between the bead images and a 2D Gaussian fit to the bead images at each z-plane. We then plotted the average localization precisions at different z-planes by the three methods against the PSF deviation values. As shown in Fig. 4, we observed significant correlations between the localization precision and the Gaussian deviation value for the quadratic and B-spline based LS fitting methods — the larger the deviation, the worse the localization precision. In contrast, WLE showed no correlation, and the determined localization precision stayed essentially flat across the different Gaussian Deviation values (Fig. 4, blue). The quadratic method sometimes even showed low precisions at low deviation PSF conditions (PSF I), which resulted from the significant deviation of the *σ_X_*(*Z*_0_), *σ_Y_*(*Z*_0_) from the quadratic calibration function for PSF I. In particular, we observed the largest Gaussian Deviation values (> 110) for PSF III, for which the corresponding localization precisions determined by the B-spline and quadratic methods degraded dramatically, whereas the WLE error remained approximately constant. In contrast, when we applied WLE and B-spline to an ideal elliptical 2D-Gaussian PSF, we obtained equally good localization precisions for both methods across different SNR conditions (Fig. S5, Supporting Material). These results strongly suggested that the distortion of PSF shape from the ideal 2D-Gaussian caused by imperfect optical setups was a major factor leading to the low localization precisions in LS fitting-based localization, and that the WLE-based localization is independent of the shape of PSF.

**Figure 4.**
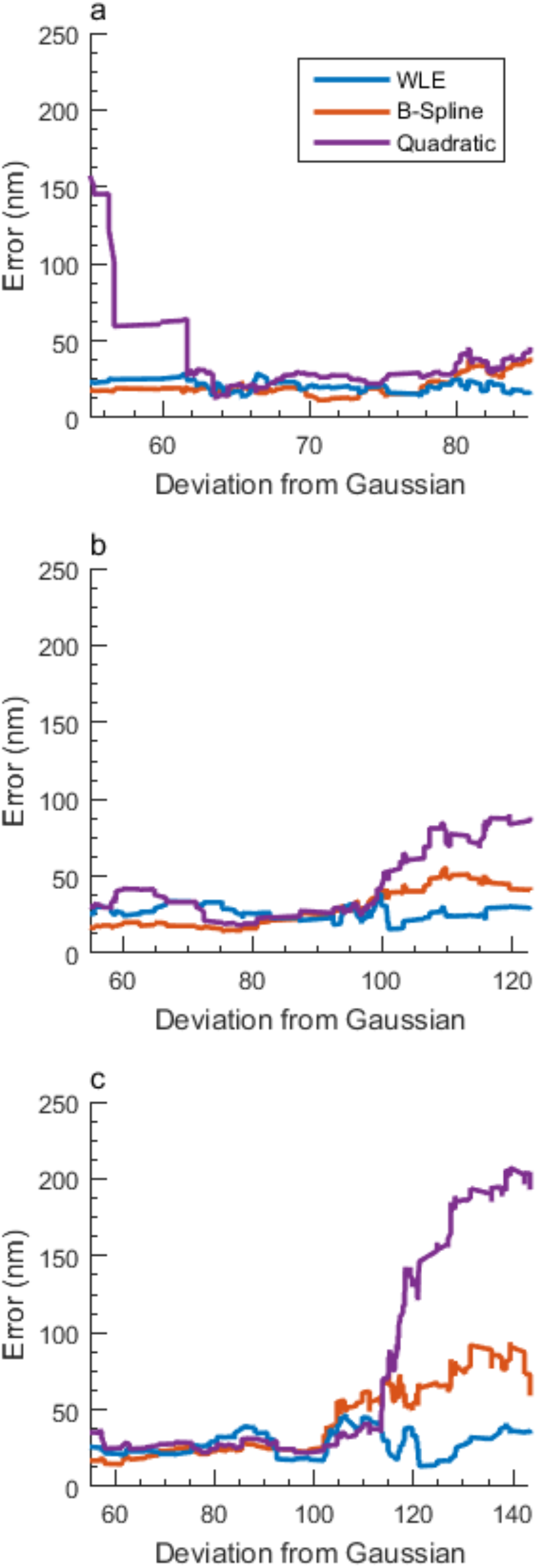
The average localization precision of LS-fitting (purple for quadratic and orange for B-spline) and WLE (blue) at different deviations of the PSF from a 2D-Gassian model.

## Discussion

In this work, we developed a WLE methodology to enhance the localization precision along the *z*-axis in astigmatism-based SMLM without an analytical description of the PSF. We validated WLE by analyzing simulated data using three experimentally measured PSFs. We found that WLE resulted in similar localization precision when compared to the commonly used B-spline fitting method when the PSF was approximately Gaussian. WLE surpassed B-spline significantly when the PSF deviated from the ideal shape, which is likely the case for real-world astigmatism-based SMLM experiments.

The major advantage of the WLE method is that it does not require a specific predefined PSF model or a user-defined noise distribution model. Both can be experimentally measured and utilized to generate calibration images, which allows WLE to predict an emitter’s z-position by numerically “matching” its image with weighted pixels to calibration images. WLE does not require the calculation of a complex pupil function, which facilitates its use by non optics-oriented users. Because of its independence of PSF shape, WLE can also be applied to BP or other PSF engineering based 3D SMLM methods. An additional novel aspect of WLE is to determine the importance of the information in each pixel (the weight) in an unbiased manner by minimizing the *z*-distance difference between repeated localizations of the same emitters.

The current WLE algorithm does not correct the PSF change caused by the refractive index mismatch. Using an experimentally measured PSF in different z-planes away from the cover glass such as fluorescence beads on another inclined surface would solve this problem(31).

The major disadvantage of WLE is that it is computationally intensive because each emitter’s image needs to be analyzed independently. Currently, the construction of a 3D superresolution image of ~2000 molecules takes ~150 CPU hours, but the current code can be further optimized for speed. With parallel or GPU computation we foresee significant improvement of the computational speed of WLE.

## Acknowledgments

This research project was conducted using computational resources at the Maryland Advanced Research Computing Center (MARCC).

## Reference

1. Hess, S.T., T.P.K. Girirajan, and M.D. Mason. 2006. Ultra-high resolution imaging by fluorescence photoactivation localization microscopy. Biophys J. 91: 4258–4272.

2. Betzig, E., G.H. Patterson, R. Sougrat, O.W. Lindwasser, S. Olenych, J.S. Bonifacino, M.W. Davidson, J. Lippincott-Schwartz, and H.F. Hess. 2006. Imaging intracellular fluorescent proteins at nanometer resolution. Science. 313: 1642–1645.

3. Rust, M.J., M. Bates, and X. Zhuang. 2006. Sub-diffraction-limit imaging by stochastic optical reconstruction microscopy (STORM). Nat Methods. 3: 793–795.

4. Baddeley, D., I.D. Jayasinghe, C. Cremer, M.B. Cannell, and C. Soeller. 2009. Light-induced dark states of organic fluochromes enable 30 nm resolution imaging in standard media. Biophys J. 96: L22–4.

5. Fu, G., T. Huang, J. Buss, C. Coltharp, Z. Hensel, and J. Xiao. 2010. In vivo structure of the E. coli FtsZ-ring revealed by photoactivated localization microscopy (PALM). PLoS ONE. 5: e12682.

6. Szymborska, A., A. de Marco, N. Daigle, V.C. Cordes, J.A.G. Briggs, and J. Ellenberg. 2013. Nuclear pore scaffold structure analyzed by super-resolution microscopy and particle averaging. Science. 341: 655–658.

7. Xu, K., G. Zhong, and X. Zhuang. 2013. Actin, spectrin, and associated proteins form a periodic cytoskeletal structure in axons. Science 339: 452–456.

8. Xiao, J., and Y.F. Dufrêne. 2016. Optical and force nanoscopy in microbiology. Nat Microbiol. 1: 16186.

9. Shroff, H., C.G. Galbraith, J.A. Galbraith, and E. Betzig. 2008. Live-cell photoactivated localization microscopy of nanoscale adhesion dynamics. Nat Methods. 5: 417–423.

10. Coltharp, C., X. Yang, and J. Xiao. 2014. Quantitative analysis of single-molecule superresolution images. Curr Opin Struct Biol. 28: 112–121.

11. Small, A., and S. Stahlheber. 2014. Fluorophore localization algorithms for super-resolution microscopy. Nat Methods. 11: 267–279.

12. Coles, B.C., S. Webb, and N. Schwartz. 2016. Characterisation of the effects of optical aberrations in single molecule techniques. Biomed Opt Exp, 7(5), 1755–1767.

13. Carlini, L., Manley, S., Carlini, L., and Manley, S. 2015. Correction of a Depth-Dependent Lateral Distortion in 3D Super-Resolution Imaging. PLoS ONE. 10: e0142949.

14. Diezmann, von, A., Y. Shechtman, and W.E. Moerner. 2017. Three-Dimensional Localization of Single Molecules for Super-Resolution Imaging and Single-Particle Tracking. Chem Rev. 117: 7244–7275.

15. Shtengel, G., J.A. Galbraith, C.G. Galbraith, J. Lippincott-Schwartz, J.M. Gillette, S. Manley, R. Sougrat, C.M. Waterman, P. Kanchanawong, M.W. Davidson, R.D. Fetter, and H.F. Hess. 2009. Interferometric fluorescent super-resolution microscopy resolves 3D cellular ultrastructure. Proc Natl Acad Sci USA. 106: 3125–3130.

16. Huang, B., W. Wang, M. Bates, and X. Zhuang. 2008. Three-dimensional super-resolution imaging by stochastic optical reconstruction microscopy. Science. 319: 810–813.

17. Pavani, S.R.P., M.A. Thompson, J.S. Biteen, S.J. Lord, N. Liu, R.J. Twieg, R. Piestun, and W.E. Moerner. 2009. Three-dimensional, single-molecule fluorescence imaging beyond the diffraction limit by using a double-helix point spread function. Proc Natl Acad Sci USA. 106: 2995–2999.

18. Ram, S., P. Prabhat, J. Chao, E.S. Ward, and R.J. Ober. 2008. High accuracy 3D quantum dot tracking with multifocal plane microscopy for the study of fast intracellular dynamics in live cells. Biophys J. 95: 6025–6043.

19. Brown, T.A., A.N. Tkachuk, G. Shtengel, B.G. Kopek, D.F. Bogenhagen, H.F. Hess, and D.A. Clayton. 2011. Superresolution fluorescence imaging of mitochondrial nucleoids reveals their spatial range, limits, and membrane interaction. Mol Cell Biol. 31: 4994–5010.

20. Lyu, Z., C. Coltharp, X. Yang, and J. Xiao. 2016. Influence of FtsZ GTPase activity and concentration on nanoscale Z-ring structure in vivo revealed by three-dimensional Superresolution imaging. Biopolymers. 105: 725–734.

21. Holtzer, L., T. Meckel, and T. Schmidt. 2007. Nanometric three-dimensional tracking of individual quantum dots in cells. Appl Phys Lett. 90(5), 053902.

22. Ovesný, M., P. Křížek, J. Borkovec, and Z. Švindrych. 2014. ThunderSTORM: a comprehensive ImageJ plug-in for PALM and STORM data analysis and super-resolution imaging. Bioinformatics, 30(16), 2389–2390.

23. Proppert, S., S. Wolter, T. Holm, and T. Klein. 2014. Cubic B-spline calibration for 3D super-resolution measurements using astigmatic imaging. Opt Express. 22(9), 10304–10316.

24. Shaevitz J.W. 2009. Bayesian estimation of the axial position in astigmatism-based threedimensional particle tracking. International Journal of Optics, 2009, Article ID 896208

25. Liu, S., E.B. Kromann, W.D. Krueger, and J. Bewersdorf. 2013. Three dimensional single molecule localization using a phase retrieved pupil function. Opt Express. 21(24), 29462–29487.

26. Petrov, P.N., Y. Shechtman, and W.E. Moerner. 2017. Measurement-based estimation of global pupil functions in 3D localization microscopy. Opt Express. 25: 7945–7959.

27. Quirin, S., S.R.P. Pavani, and R. Piestun. 2011. Optimal 3D single-molecule localization for superresolution microscopy with aberrations and engineered point spread functions. Proc Natl Acad Sci USA. 109: 675–679.

28. Abraham, A.V., S. Ram, J. Chao, E.S. Ward, and R.J. Ober. 2010. Quantitative study of single molecule location estimation techniques. Opt Express. 17: 23352–23373.

29. Ober, R.J., S. Ram, and E.S. Ward. 2004. Localization accuracy in single-molecule microscopy. Biophys J. 86: 1185–1200.

30. Izeddin, I., J. Boulanger, V. Racine, C.G. Specht, A. Kechkar, D. Nair, A. Triller, D. Choquet, M. Dahan, and J.B. Sibarita. 2012. Wavelet analysis for single molecule localization microscopy. Opt Express. 20: 2081–2095.

31. Besseling, T.H., J. Jose, and A. Van Blaaderen. 2014. Methods to calibrate and scale axial distances in confocal microscopy as a function of refractive index. J Microsc. 257: 142–150.

